# Effect of seminal plasma extracellular vesicles in post-thaw functional parameters of cryopreserved ram sperm

**DOI:** 10.64898/2026.06.17.732841

**Authors:** A.R. Nicolli, T. Armani, M. Buendía Arellano, L. Zalazar, F. A. Hozbor, A. Cesari

## Abstract

Cryopreservation of ram semen induces structural and functional alterations that compromise sperm function. Given that seminal plasma contributes to sperm physiology regulation, increasing attention has focused on extracellular vesicles (EVs), which carry proteins involved in membrane organization and capacitation. These properties suggest that EVs may contribute to maintaining sperm stability during cryopreservation. Therefore, this study evaluated the effect of seminal plasma-derived EVs on post-thaw functional parameters of ram sperm. Semen was cryopreserved in the presence or absence of EVs isolated by ultracentrifugation previously characterized by nanoparticle tracking analysis, TEM, FTIR and Western blotting. Post-thaw sperm quality was assessed through viability, membrane lipid disorder, reactive oxygen species production, protein phosphorylation, acrosome status, intracellular calcium levels, and motility. Sperm cryopreserved with EV-supplemented extender showed reduced membrane lipid disorder and lower intracellular calcium levels compared to controls (p < 0.05). CASA analysis revealed no significant differences in total or progressive motility; however, EV supplementation was associated with a modified kinematic profile, characterized by increased velocity and trajectory-related parameters (p < 0.001), suggesting a potential effect on sperm movement patterns. No significant differences were observed in viability, ROS levels, protein phosphorylation (tyrosine residues or PKA), or acrosome status. These findings provide proof-of-concept evidence that supplementation with seminal plasma-derived EVs during ram semen cryopreservation is associated with changes in membrane lipid organization, intracellular calcium homeostasis, and selected sperm kinematic parameters. Further studies are required to determine whether these changes translate into improved fertilizing ability.

**Highlights:**

- Seminal EVs modulates post-thaw functional parameters in ram sperm.
- EVs reduce membrane lipid disorder and intracellular Ca²⁺ levels.
- EVs modify sperm kinematics, increasing linearity and straightness.
- EVs do not affect viability, ROS, phosphorylation or acrosome status.
- EV supplementation provides proof-of-concept for improving selected post-thaw sperm functions.

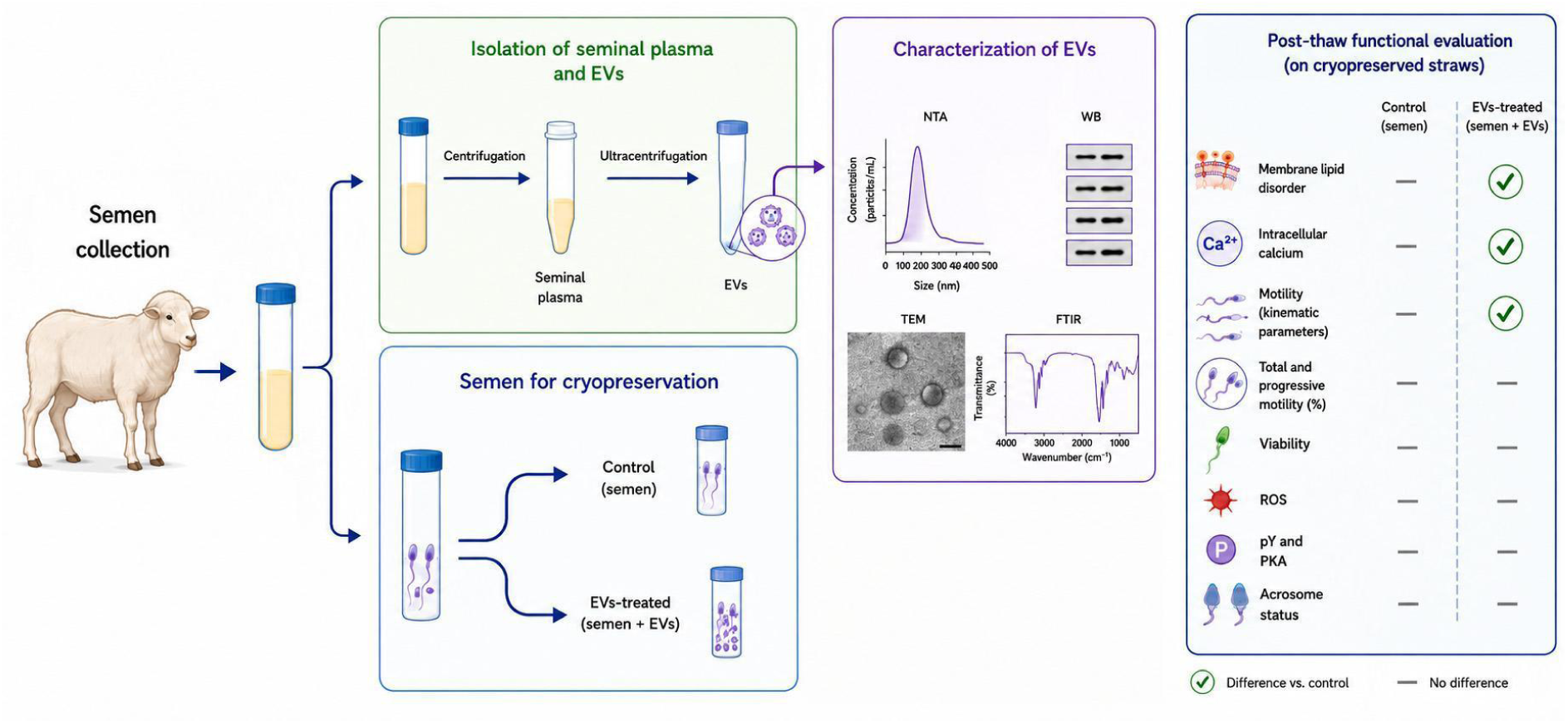

## 1. Introduction

Semen cryopreservation is considered an important tool for animal breeding and genetic conservation programs, as it allows the long-term preservation of genetic material for potential use in artificial insemination. However, despite continuous improvements in freezing protocols and extender formulations, post-thaw sperm quality in rams remains highly variable and often suboptimal when compared with other domestic species (Salamon and Maxwell 2000; Watson 2000; Gillan et al. 2004). The freeze-thaw process induces structural and functional alterations collectively referred to as cryoinjury, caused by thermal and osmotic shocks, dehydration shock, ice crystal formation and elevated levels of reactive oxygen species (ROS) (Stanic et al. 2000; Thomson et al. 2009; Yeste 2015; Valipour et al. 2021), which compromise sperm viability and fertilizing potential. In ram sperm, cryopreservation has been associated with reduced motility, altered membrane integrity, premature capacitation-like changes, and impaired sperm functionality after thawing (Bailey et al. 2000; Watson 2000; Martínez-Pastor et al. 2010). These limitations highlight the need for novel, biologically relevant strategies to mitigate cryodamage and improve post-thaw sperm quality in this species.

The sperm plasma membrane is one of the primary targets of cryoinjury due to its unique lipid composition and high degree of organization, which are essential for sperm function. In ram spermatozoa, freezing and thawing disrupt membrane lipid architecture, leading to increased membrane fluidity, loss of lipid asymmetry, and altered permeability (Bailey et al. 2000; Watson 2000). These changes can be detected as an increase in membrane lipid disorder, which has been associated with reduced sperm viability and functional competence (Rathi et al. 2001). Membrane destabilization during cryopreservation is closely linked to dysregulation of intracellular calcium (Ca²⁺) homeostasis. Increased membrane permeability facilitates uncontrolled Ca²⁺ influx, triggering premature capacitation-like events, altered motility patterns, and early activation processes that compromise sperm fertilizing ability (Bailey et al. 2000; Martínez-Pastor et al. 2010). Therefore, preservation of membrane lipid organization and regulation of intracellular Ca²⁺ levels are critical determinants of post-thaw sperm quality.

Seminal plasma is a complex and biologically active fluid that plays a crucial role in regulating sperm physiology beyond serving as a transport medium. It contains a wide variety of proteins, lipids, enzymes, and signaling molecules that modulate sperm membrane stability, motility, capacitation, and survival within the female reproductive tract (Druart et al. 2013; Maxwell et al. 2007). In rams, seminal plasma components have been shown to interact directly with fresh and cryopreserved spermatozoa, influencing membrane composition and functional status (Barrios et al. 2000; Bernardini et al. 2011; Ramírez-Vasquez et al. 2019). Although supplementation with whole seminal plasma has shown beneficial effects on sperm function under certain experimental conditions, its application during cryopreservation remains controversial because seminal plasma also contains factors that may impair sperm preservation, including proteins associated with membrane destabilization, variability among males and ejaculates, and components that may promote premature capacitation-like changes or reduce cryosurvival when present at inappropriate concentrations (Bernardini et al. 2011; Ledesma et al. 2016; Ramírez-Vasquez et al. 2019). Therefore, identifying specific bioactive fractions responsible for the beneficial effects of seminal plasma may provide a more consistent and targeted strategy to improve cryopreservation outcomes. Among the bioactive components of seminal plasma, extracellular vesicles (EVs) have emerged as key mediators of cell-to-cell communication in the male reproductive tract. Seminal plasma-derived EVs are abundant and heterogeneous membrane-enclosed nanovesicles that carry proteins, lipids, and nucleic acids capable of modulating target cell function (Vojtech et al. 2014; Simon et al. 2018a; Leahy et al. 2020). Several studies have demonstrated that seminal plasma EVs can bind to spermatozoa and transfer bioactive cargo, thereby influencing sperm maturation, membrane remodeling, calcium signaling, capacitation, acrosome reaction, and motility (Sullivan and Saez 2013; Aalberts et al. 2014). Because EVs are natural lipid-based carriers, their phospholipid bilayer protects their molecular cargo from degradation and facilitates its efficient delivery and fusion with the sperm plasma membrane, enabling the transfer of bioactive lipids, proteins and regulatory RNAs involved in membrane remodeling and cell signaling (Sullivan and Saez 2013; Aalberts et al. 2014; Armani et al. 2026). In ovine species, seminal plasma EVs have been shown to interact with specific regions of the sperm cell and carry proteins potentially involved in the regulation of sperm functional competence (Barranco et al. 2023; Armani et al. 2026). These findings suggest that EVs may play a physiological role in maintaining sperm functionality.

To date, no studies have evaluated the effect of EV supplementation of the extender used for cryopreservation of ram semen, nor its impact on post-thaw membrane organization, intracellular calcium regulation, or sperm motility. Therefore, the aim of the present study was to evaluate, for the first time, the effect of seminal plasma-derived extracellular vesicles added during ram semen cryopreservation on post-thaw functional parameters. The central hypothesis of this study was that seminal plasma-derived extracellular vesicles (EVs) protect spermatozoa against cryoinjury by preserving plasma membrane integrity, thereby reducing membrane destabilization, limiting premature capacitation-like changes, maintaining intracellular Ca²⁺ homeostasis, and improving post-thaw sperm functionality.

## 2. Materials and Methods

### 2.1 Animals, semen collection and cryopreservation

Semen was collected from nine mature Texel rams maintained at the Experimental Station of Balcarce (Instituto Nacional de Tecnología Agropecuaria, INTA, Argentina) during the natural breeding season (autumn). All animal procedures were performed in accordance with institutional guidelines for animal care and were approved by the Animal Ethics Committee of INTA (CICUAE INTA CeRBAS; protocol ID 217/2021).

Ejaculates were collected using an artificial vagina and evaluated immediately after collection. Only ejaculates exhibiting a mass motility score ≥ 4 (scale 1-5) were included in the study. Cryopreservation experiments were conducted in three separate sessions during the breeding season. In each session, a pooled semen sample was prepared by combining ejaculates from three rams randomly selected from the nine rams included in the study that met the quality criteria described above. These semen pools were different from those used for seminal plasma collection and EV isolation.

Each pooled semen sample was diluted to a final concentration of 400 × 10⁶ spermatozoa/mL at 37°C using a commercial soy lecithin-based extender (AndroMed®, Minitube, Germany). The diluted semen was divided into two equal aliquots corresponding to the following experimental groups: (i) Control, without extracellular vesicles (EVs), and (ii) EV-treated, supplemented with seminal plasma-derived EVs. Based on the supplementation strategy previously described by Zhang et al. (2024), EVs were resuspended in diluted AndroMed® extender and added at a final volume corresponding to approximately 10% of the initial semen volume, with the aim of supplementing the extender while minimizing changes in its composition. According to the particle concentration of each EV preparation, determined by nanoparticle tracking analysis, this volume corresponded to approximately 1.3 × 10¹³ EV particles per biological replicate. Preliminary assays confirmed that replacing 10% of the extender volume with EV suspension did not adversely affect sperm viability. A different EV preparation, obtained from one of the three seminal plasma pools and previously characterized, was used in each cryopreservation session. Consequently, three biological replicates were generated, each consisting of a distinct pooled semen sample evaluated in the presence or absence of a different EV preparation.

Samples were cooled from 30°C to 7°C over 2 h and maintained at 7°C for an additional 1.5 h before packaging into 0.25-mL plastic straws containing 0.2 mL of semen. Straws were sealed with polyvinyl alcohol powder and frozen by exposure to liquid nitrogen vapour (5 cm above the liquid nitrogen surface for 10 min) before being plunged into liquid nitrogen for long-term storage.

For post-thaw analyses, semen straws were thawed in a water bath at 37°C for 30 s and layered onto 400 µL of OviPure® colloid (Minitube, Germany). Sperm selection was performed according to the manufacturer’s instructions (Morrell and Jasim, 2021). Briefly, samples were centrifuged at 800 × g for 10 min to remove seminal plasma and non-viable spermatozoa. The recovered sperm pellet was washed once in phosphate-buffered saline (PBS; pH 7.4) by centrifugation at 800 × g for 10 min and finally resuspended to the concentration required for each subsequent analysis.

### 2.2 Seminal plasma (SP) preparation

Seminal plasma used for EV isolation was collected during three independent sampling sessions throughout the autumn breeding season. In each session, ejaculates from three mature Texel rams meeting the quality criteria described above were pooled to generate one seminal plasma sample. Thus, three independent seminal plasma pools were obtained, each processed independently for EV isolation and characterization. For seminal plasma (SP) preparation, the same semen quality criteria described above were applied. Fresh pooled ejaculates obtained from three rams were centrifuged at 800 × g for 10 min at room temperature to pellet spermatozoa. The resulting supernatant was subsequently centrifuged twice at 16,000 × g for 30 min at 4°C to eliminate residual cells, cell debris and large particulate material. The resulting SP was examined microscopically to confirm the absence of spermatozoa before being used for extracellular vesicle (EV) isolation and stored at −80°C until use.

### 2.3 Isolation and characterization of seminal plasma extracellular vesicles (EVs)

Extracellular vesicles (EVs) were isolated by differential ultracentrifugation. Briefly, 700 μL of SP were diluted with filtered phosphate-buffered saline (PBS) to a final volume of 3.2 mL and centrifuged twice at 100,000 × g for 1.5 h at 4°C. The resulting EV pellet was resuspended in diluted AndroMed® extender (1:5, v/v; one part extender to five parts water for injection) and stored at 4°C until use.

Protein concentration was determined using a NanoDrop One spectrophotometer (Thermo Scientific, USA) with bovine serum albumin (BSA) as the calibration standard. EV preparations were subsequently characterized by nanoparticle tracking analysis (NTA) and Western blotting following the Minimal Information for Studies of Extracellular Vesicles (MISEV 2023) guidelines (Welsh et al. 2024). Each EV preparation was characterized for particle concentration and protein markers before being used in the corresponding cryopreservation experiment.

### 2.4 Transmission electron microscopy (TEM) and Fourier-transform infrared spectroscopy (FTIR)

For transmission electron microscopy, isolated EVs were processed following a modification of the procedure described by Théry et al. (2006). EV pellets were resuspended in 2% w/v paraformaldehyde (PFA) in PBS and incubated overnight at 4°C. Then, 10 μL of the EV preparation were deposited onto formvar-coated TEM grids and allowed to adsorb for 20 min. Samples were post-fixed in 1% w/v glutaraldehyde, washed with distilled water, and contrasted with uranyl oxalate (pH 7), followed by methyl cellulose-UA (pH 4) on ice for negative contrasting. EVs were visualized using a Hitachi HT 7800 transmission electron microscope (Hitachi, Tokyo, Japan) operated at 80 kV and photographed with an NS 15 AMT camera (Advanced Microscopy Techniques Corp., Woburn, MA, USA). TEM imaging was performed at the Transmission Electron Microscopy Service of the Instituto de Investigaciones en Ciencias de la Salud (INICSA)-CONICET, Centro de Microscopía Electrónica, Facultad de Ciencias Médicas, Universidad Nacional de Córdoba, Argentina.

For Fourier-transform infrared spectroscopy (FTIR), the composition of the EV-enriched fraction was characterized to confirm its biological origin. Spectra were collected by attenuated total reflectance (ATR) using a Nicolet 6700 (Thermo Scientific, USA) spectrometer fitted with a diamond-crystal ATR accessory at INTEMA-CONICET. All measurements were performed at room temperature under ambient conditions. For each sample, 25 μL were deposited onto the 2-mm-diameter sampling area of the ATR crystal and allowed to air-dry before analysis. A total of 128 scans were co-added at a nominal resolution of 4 cm⁻¹ over a spectral range of 4000-400 cm⁻¹.

### 2.5 Nanoparticle tracking analysis (NTA)

Particle concentration and size distribution were determined by nanoparticle tracking analysis (NTA) using a ZetaView® Twin Laser instrument (PMX-230, Particle Metrix, Germany) operated at 25°C. Three independent EV preparations (one obtained from each pooled seminal plasma sample) were analyzed. Each preparation was measured in eleven consecutive acquisitions, and the average value was used for subsequent analyses. Data acquisition and processing were performed using the manufacturer’s software according to the MISEV 2023 recommendations (Welsh et al. 2024). NTA analyses were carried out at the Instituto de Investigaciones en Microbiología y Parasitología Médica (UBA-CONICET, Argentina).

### 2.6 Western blot analysis

Western blot analysis was performed using three independent EV preparations. Equal amounts of EV proteins were denatured, separated by electrophoresis on 15% SDS-polyacrylamide gels and transferred onto polyvinylidene difluoride (PVDF) membranes (Bio-Rad, USA).

Membranes were blocked for 1 h at room temperature with 3% (w/v) skimmed milk prepared in Tris-buffered saline containing 0.05% Tween-20 (TBS-T; 10 mM Tris-HCl, pH 8.0, 120 mM NaCl) and incubated with the following primary antibodies: anti-CD9 (1:1000; D3H4P, Cell Signaling Technology), anti-HSP70 (1:2000; SAB4200714, Sigma-Aldrich), anti-calnexin (1:1000; ab22595, Abcam) and anti-ApoC3 (1:3000; PA5-114865, Invitrogen). Membranes were then incubated with horseradish peroxidase-conjugated anti-mouse or anti-rabbit IgG secondary antibodies (1:10,000; Sigma-Aldrich). Between successive immunodetections, membranes were stripped using stripping buffer (60 mM Tris-HCl, pH 6.8, 2% SDS and 0.8% β-mercaptoethanol), washed extensively and re-blocked before incubation with the next antibody. Immunoreactive bands were visualized using enhanced chemiluminescence (ECL Plus, GE Healthcare) and detected with a LI-COR C-DiGit chemiluminescence scanner.

### 2.7 Post-thaw sperm evaluation

Frozen semen straws were thawed as described above and maintained at 37°C throughout all analyses. Following OviPure® selection, sperm concentration was determined using a counting chamber and adjusted according to the requirements of each assay. Unless otherwise indicated, all post-thaw evaluations were performed using three independent pooled semen samples (biological replicates), each prepared from ejaculates of three rams and analyzed under both Control and EV-treated conditions. Sperm motility was initially assessed by phase-contrast microscopy to evaluate post-thaw sample quality. Only samples exhibiting >80% subjective motility were included in subsequent analyses. For flow cytometry assays, sperm concentration was adjusted to 80-100 × 10⁶ spermatozoa/mL in phosphate-buffered saline (PBS, pH 7.4), unless otherwise specified. For each analysis, 15 μL of sperm suspension (approximately 1.2-1.5 × 10⁶ spermatozoa) were used.

Intracellular reactive oxygen species (ROS) production was evaluated using 2’,7’-dichlorodihydrofluorescein diacetate (DCFDA; 25 μM; Invitrogen, C6827). Samples were incubated for 20 min at 37°C in the dark and analyzed immediately by flow cytometry. ROS levels were expressed as the mean DCF fluorescence intensity (arbitrary units, AU). Sperm viability was assessed using propidium iodide (PI; 1 μg/mL; Sigma-Aldrich, P4170), which was added immediately before acquisition. Viable spermatozoa were identified as PI-negative cells.

Acrosomal status was evaluated using peanut agglutinin conjugated to Alexa Fluor 488 (PNA-Alexa Fluor 488; 1 μg/mL). Sperm samples were incubated for 10 min at 37°C in the dark before analysis. Acrosome-reacted spermatozoa were identified as PNA-positive/PI-negative (PNA+/PI-) cells. Sperm treated with the calcium ionophore A23187 (10 μM for 10 min at 37°C) were included as positive controls.

Intracellular calcium levels were determined using the fluorescent probe Fluo-3 AM (10 μM; Invitrogen, F1242). For this assay, sperm concentration was adjusted to 80–100 × 10⁶ spermatozoa/mL using synthetic oviductal fluid (SOF; 107.70 mM NaCl, 7.17 mM KCl, 1.19 mM KH₂PO₄, 1.71 mM CaCl₂, 0.49 mM MgCl₂, 25.07 mM NaHCO₃, 3.30 mM sodium lactate, 0.30 mM sodium pyruvate and 20 mM HEPES), which provides physiological ionic conditions for calcium measurements. Samples were incubated with Fluo-3 AM for 20 min at 37°C in the dark and analyzed by flow cytometry. Positive controls consisted of spermatozoa treated with A23187 (10 μM for 15 min at 37°C). Intracellular calcium levels were expressed as the percentage of Fluo-3-positive spermatozoa.

Membrane lipid disorder was evaluated using Merocyanine 540 (M540). Sperm concentration was adjusted to 10 × 10⁶ spermatozoa/mL and cells were incubated with M540 (1.5 μM; Sigma-Aldrich, 323756) for 10 min at 37°C in the dark. Yo-Pro-1 iodide (2.25 μM) was subsequently added to identify non-viable spermatozoa. Fluorescence was evaluated using a Nikon C1SiR confocal microscope under identical acquisition settings for all experimental groups. Membrane lipid disorder was quantified as the percentage of M540-positive/Yo-Pro-1-negative spermatozoa. Sperm treated with 1% Triton X-100 followed by freezing at −20°C were included as positive controls for membrane disruption.

For protein phosphorylation analyses, spermatozoa were collected by centrifugation at 10000 × g for 10 min. Pellets were resuspended in 30 μL of 5× sample buffer, vortexed for 30 s and boiled for 5 min. Samples were stored overnight at −20°C, supplemented with β-mercaptoethanol (20%, v/v), boiled again for 5 min and loaded onto 12% SDS-polyacrylamide gels (2 × 10⁶ spermatozoa per lane). Western blotting was performed as described above using anti-phosphotyrosine (1:1000; clone 4G10, Millipore), anti-PKA substrate (1:5000; clone 100G7E, Cell Signaling Technology) and anti-α-tubulin (1:10000; Sigma-Aldrich, T6074) antibodies. Between successive immunodetections, membranes were stripped, washed and re-blocked as described above.

### 2.8 Sperm motility assessment

Sperm motility was evaluated using a computer-assisted sperm analysis (CASA) system. Analyses were performed using the same three independent pooled semen samples (biological replicates) described above, each evaluated under Control and EV-treated conditions. Following OviPure® selection, sperm concentration was adjusted to 50 × 10⁶ spermatozoa/mL in PBS supplemented with 5% fructose. OviPure® selection was performed to recover the motile sperm population and remove cryoprotectant residues, debris, and non-viable spermatozoa prior to CASA analysis. Since both Control and EV-treated samples underwent the same procedure, any effect of sperm selection was consistent across treatments. A 7-μL aliquot was loaded into a 10-μm Cell-Vu chamber (Millennium Sciences, USA) and examined under a Nikon Eclipse E200 microscope equipped with a 20× negative phase-contrast objective. Video recordings were acquired using a digital camera set at 600 TV-line resolution, with autofocus disabled, at 30 frames/s. Three randomly selected microscopic fields were recorded per biological replicate, at least 200 spermatozoa were analyzed per sample.

Objective motility analysis was performed using the CASA system following the recommendations of Wang et al. (2020) and based on the analytical approach described by Buchelly Imbachi et al., (2018). The following kinematic parameters were recorded: curvilinear velocity (VCL, μm/s), straight-line velocity (VSL, μm/s), average path velocity (VAP, μm/s), linearity (LIN, %), straightness (STR, %), wobble (WOB, %), amplitude of lateral head displacement (ALH, μm), beat-cross frequency (BCF, Hz), total motility (TM, %) and progressive motility (PM, %), the latter defined as spermatozoa exhibiting STR ≥80%.

### 2.9 Statistical analysis

All statistical analyses were performed using R software version 4.1.3 (R Foundation for Statistical Computing, Vienna, Austria). Data were analyzed using mixed-effects models to account for the experimental structure, including biological replicate as a random effect and treatment as a fixed effect. Continuous variables were analyzed using linear mixed-effects models (LMMs). These analyses included sperm viability, total and progressive motility, ROS production, intracellular calcium levels, phosphorylation levels of PKA and tyrosine residues, and CASA-derived kinematic parameters. For CASA analysis, individual sperm measurements were included as observations, while biological replicate was incorporated as a random effect to account for measurements obtained from multiple spermatozoa within each independent experiment. Variables representing proportions or bounded continuous data were analyzed using generalized linear mixed-effects models (GLMMs) with appropriate distributions according to data characteristics. Membrane lipid disorder assessed by merocyanine 540 staining was analyzed using a beta-distributed GLMM. Model assumptions were evaluated by inspection of residual distributions and diagnostic plots. Estimated marginal means and treatment contrasts were obtained using the emmeans package. Results are reported as estimated treatment effects, 95% confidence intervals, and p-values. Statistical significance was defined as p < 0.05.

## 3. Results

### 3.1 Extracellular vesicle characterization

Extracellular vesicles (EVs) were isolated by ultracentrifugation (UC) from ram seminal plasma and subsequently characterized by nanoparticle tracking analysis (NTA), transmission electron microscopy (TEM), Western blotting, and attenuated total reflectance Fourier-transform infrared spectroscopy (ATR-FTIR). NTA revealed a heterogeneous population of particles with a major size peak centered around 100-120 nm in diameter (**Figure 1a**). Particle size analysis showed a mean diameter of 120.30 ± 2.43 nm and a median (X50) of 105.70 ± 0.92 nm, with X10 and X90 values of 68.37 ± 0.75 nm and 179.93 ± 6.92 nm, respectively. The mean particle concentration was 1.43 × 10¹⁴ ± 0.98 × 10¹⁴ particles/mL. Particle size distribution was consistent across biological replicates, and measurement of particle content in the AndroMed™ extender used as a control revealed negligible levels compared with the EV preparations (**Figure 1a**). TEM images confirmed the presence of vesicular structures with the characteristic membranous bilayer morphology of EVs, consistent with the size range observed by NTA (**Figure 1b**).

**Figure 1.**
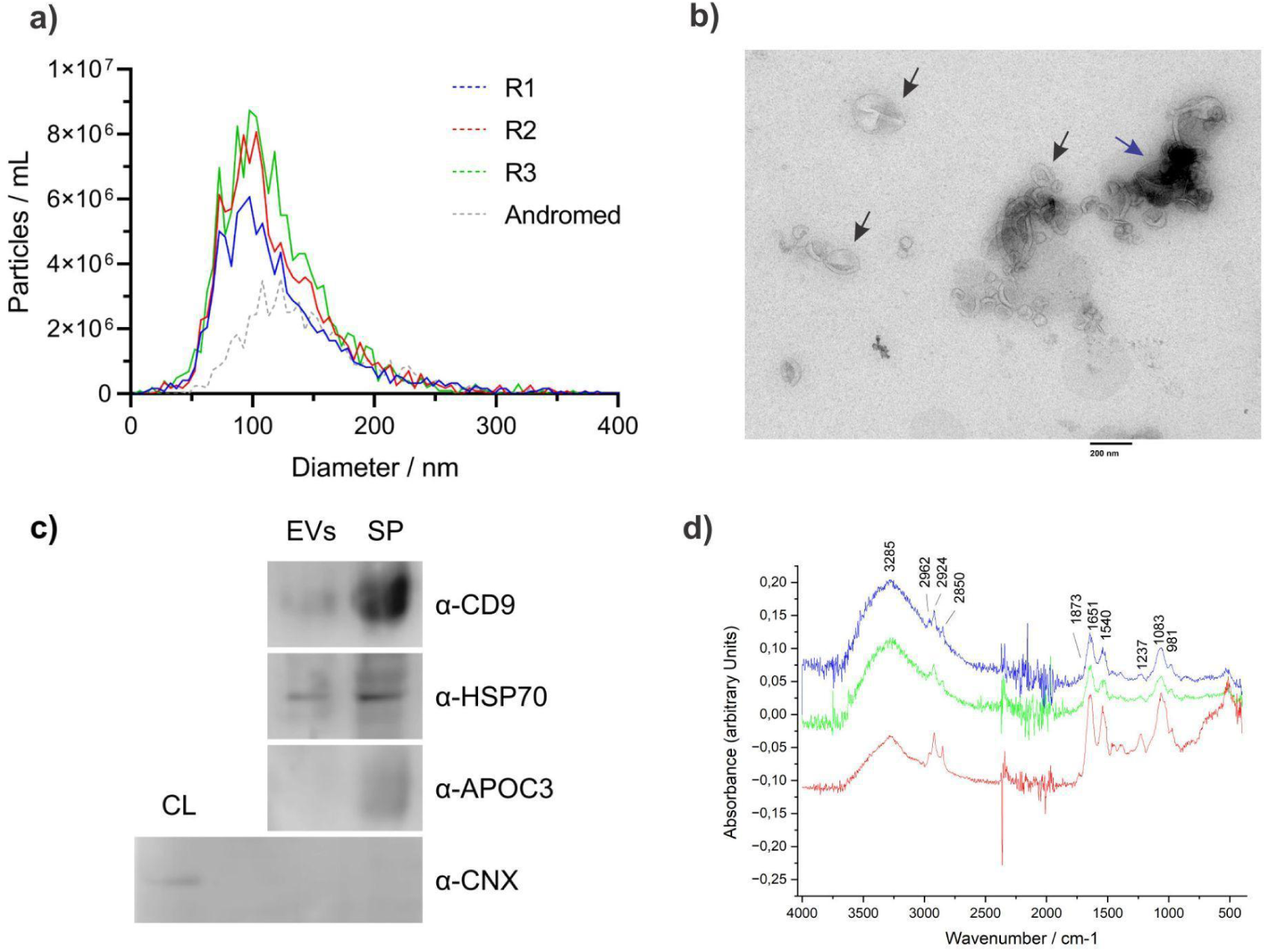
Isolation and characterization of extracellular vesicles (EVs) derived from ram seminal plasma. (a) Representative nanoparticle tracking analysis (NTA) profiles showing the size distribution and particle concentration of EVs isolated from seminal plasma across three biological replicates (R1-R3). The AndroMed™ extender was analyzed as a negative control. (b) Transmission electron microscopy (TEM) image of isolated EVs showing typical cup-shaped morphology. Black arrows indicate EVs of varying sizes, while the blue arrow indicates protein aggregates. Scale bar = 200 nm. (c) Representative Western blot analysis of EVs and seminal plasma (SP) showing the presence of EV-associated markers CD9 and HSP70 and the absence of the non-vesicular extracellular particle marker ApoC3. Calnexin (CNX), an endoplasmic reticulum marker, was detected only in the whole-cell lysate (CL, positive control), confirming antibody specificity and minimal cellular contamination. Full-length Western blots are provided in **Supplementary Figure 1**. (d) ATR-FTIR spectra of seminal plasma extracellular vesicles (EVs). Characteristic absorption bands are indicated on the plot in decreasing wavenumber order: the broad band around **3285 cm**⁻**¹**, corresponding to N–H stretching vibrations of peptide groups in proteins (amide A); the bands at **2924 and 2850 cm**⁻**¹**, assigned to the asymmetric and symmetric stretching vibrations of lipid acyl CH₂ groups, respectively; the absorption band around **1738 cm**⁻**¹**, arising from C=O stretching of ester groups in phospholipids, triglycerides, and cholesterol esters; the prominent amide I band around **1651 cm**⁻**¹**, mainly associated with C=O stretching vibrations of the protein peptide backbone; the amide II band around **1540 cm**⁻**¹**, arising primarily from N–H bending vibrations of peptide groups; the band at **1453 cm**⁻**¹**, attributed to bending vibrations of lipid acyl CH₂ groups; and the band at **1394 cm**⁻**¹**, assigned to bending vibrations of lipid and protein CH₃ groups. Bands in the **1200–950 cm**⁻**¹** region are attributed to phosphodiester stretching vibrations of phospholipids and C–O–C stretching vibrations of phospholipids, triglycerides, and cholesterol esters; however, this spectral region is masked by the broad phosphate vibration bands of the PBS buffer. Experiments were performed with three independent biological replicates (n = 3) for NTA, Western blot, TEM, and FTIR analyses; representative images are shown.

Western blot analysis further confirmed the vesicular nature of the isolated particles. EV preparations were positive for the EV-associated markers CD9 and HSP70 (**Figure 1c**). In addition, apolipoprotein C3 (ApoC3), a marker associated with non-vesicular extracellular particles (NVPs), was detected in the seminal plasma but was nearly absent in the isolated EVs enrichment fraction. Moreover, calnexin (CNX), an endoplasmic reticulum marker, was detected exclusively in the cellular lysate (CL) but was not found in the EV samples, indicating minimal contamination with intracellular and NVPs components (**Figure 1c**).

ATR-FTIR spectra of the EV-enriched fraction showed the typical molecular composition of biological samples, with characteristic absorption bands corresponding to proteins (amide I and II bands), lipids (CH₂ stretching bands), and phospholipids (phosphodiester stretching bands), further supporting the vesicular nature and purity of the isolated EVs (**Figure 1d**).

### 3.2 EV supplementation does not affect sperm viability, acrosomal status, or oxidative stress

Post-thaw sperm viability was comparable between control and EV-treated samples, indicating that the presence of extracellular vesicles during cryopreservation did not significantly modify post-thaw sperm viability (**Figure 2a; Supplementary Table 1**). Likewise, the percentage of acrosome-reacted spermatozoa (assessed by PNA-Alexa 488 staining) and the intracellular reactive oxygen species (ROS) levels, did not differ between experimental groups, suggesting that EV supplementation did not significantly affect these post-thaw parameters under the evaluated conditions (**Figure 2b,c; Supplementary Table 1**).

**Figure 2.**
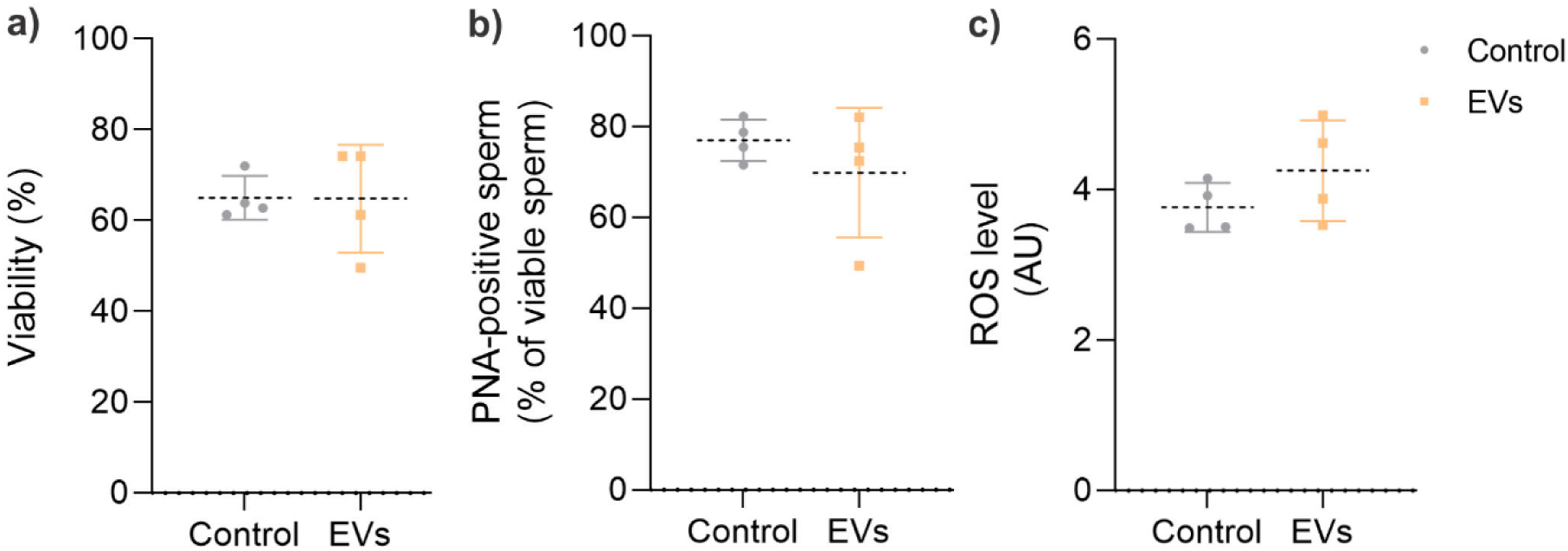
Effect of seminal plasma extracellular vesicles (EVs) on viability, acrosomal status and oxidative stress of post-thaw ram spermatozoa. Ram spermatozoa were cryopreserved in the absence (Control) or presence of EVs. Analyses were performed using three independent pooled semen samples, each prepared from ejaculates of three rams. (a) Sperm viability expressed as the percentage of viable cells (IP−/total). (b) Percentage of acrosome-reacted viable spermatozoa (PNA+/IP−). (c) Intracellular reactive oxygen species (ROS) levels determined by DCFDA fluorescence and expressed as arbitrary fluorescence units (AU). Each dot represents one biological replicate (pooled ejaculates from three rams), and dashed lines indicate the mean. Data are presented as mean ± SEM. No significant differences were detected between treatments (p > 0.05).

### 3.3 EVs supplementation is associated with reduced membrane lipid disorder and intracellular calcium levels without altering capacitation-associated signaling markers

Spermatozoa cryopreserved in the presence of EVs exhibited a significantly lower M540 signal in viable spermatozoa, indicating reduced membrane lipid disorder, compared with control samples (OR = 0.187; 95% CI: 0.109-0.322; p < 0.001;; **Figure 3a,b**). Additionally, intracellular calcium levels were significantly lower in EV-treated spermatozoa relative to controls (estimate = −5.63; 95% CI: −11.3-0.008; p = 0.048; **Figure 3c,d**). Importantly, this stabilization of membrane organization and Ca²⁺ homeostasis was not accompanied by changes in capacitation-associated signaling pathways, as no differences were observed in PKA activity or protein tyrosine phosphorylation between groups (**Figure 3e,f; Supplementary Table 1**). Together, these results indicate that EV supplementation during cryopreservation is associated with improved membrane lipid organization and lower intracellular Ca^2+^ levels, without detectable changes in downstream capacitation-associated signaling pathways in post-thaw ram spermatozoa under the conditions tested.

**Figure 3.**
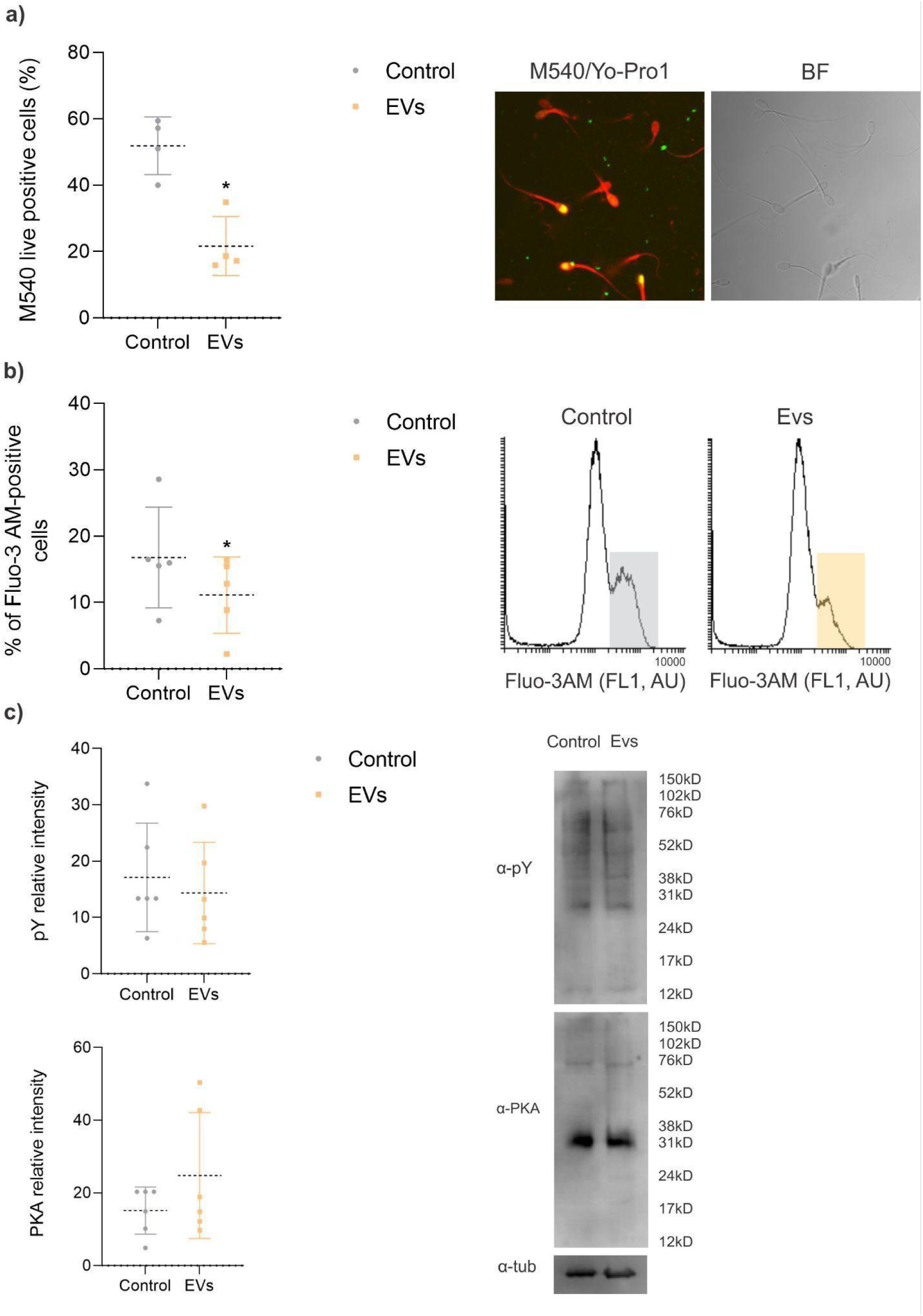
Effect of seminal plasma extracellular vesicles (EVs) on membrane lipid disorder, intracellular calcium levels and protein phosphorylation in post-thaw ram spermatozoa. Ram spermatozoa were cryopreserved in the absence (Control) or presence of EVs. Data were obtained from three independent biological replicates (pooled ejaculates from three rams per replicate). (a) Membrane lipid disorder assessed by Merocyanine 540 (M540) staining. Left, percentage of M540-positive viable spermatozoa. Right, representative confocal microscopy images showing merged fluorescence of M540 (red) and Yo-Pro-1 (green), and the corresponding bright-field (BF) images. Cells stained only with M540 appear red, whereas cells positive for both M540 and Yo-Pro-1 appear yellow. (b) Intracellular calcium levels evaluated by flow cytometry using Fluo-3 AM. Left, percentage of Fluo-3 AM-positive cells. Right, representative fluorescence histograms of Control and EV-treated spermatozoa; the vertical dashed line indicates the threshold for positive cells. (c) Protein phosphorylation analysis. Left, densitometric quantification of phosphotyrosine-containing proteins (pY) and protein kinase A substrate-phosphorylated proteins (PKA), normalized to α-tubulin. Right, representative Western blots (2 × 10⁶ spermatozoa per lane) probed with anti-phosphotyrosine (α-pY), anti-PKA substrate (α-PKA), and anti-α-tubulin (α-Tub) antibodies. Data are presented as mean ± SEM. Full-length Western blots are provided in **Supplementary Figure 2**.

### 3.4 EV supplementation modifies sperm kinematic parameters without altering motility percentages

The percentage of total and progressively motile spermatozoa was similar between the control and EV-treated groups (**Table 1; Supplementary Table 1**). However, mixed-model analysis of CASA parameters revealed significant differences in sperm kinematic profiles following EV supplementation. EV-treated spermatozoa exhibited higher velocity-related parameters, including VCL (estimate = 84.3 µm/s; 95% CI: 69.2-99.4; p < 0.001), VAP (estimate = 78.7 µm/s; 95% CI: 63.6-93.7; p < 0.001), and VSL (estimate = 77.6 µm/s; 95% CI: 62.2-92.9; p < 0.001). In addition, EV supplementation was associated with increased values of trajectory-related parameters, including LIN (estimate = 9.15; 95% CI: 6.82-11.5; p < 0.001), STR (estimate = 5.79; 95% CI: 4.06-7.53; p < 0.001), and WOB (estimate = 6.63; 95% CI: 4.88-8.38; p < 0.001). ALH and BCF were also higher in EV-treated samples (ALH: estimate = 0.783; 95% CI: 0.615-0.951; BCF: estimate = 2.60; 95% CI: 1.93-3.27; both p < 0.001). Together, these findings indicate that EV supplementation was associated with a modified post-thaw sperm kinematic profile, characterized by increased velocity-related parameters and altered trajectory characteristics, while the proportion of motile spermatozoa remained unchanged.

**Table 1.** Motility parameters of post-thaw ram spermatozoa cryopreserved in the absence (Control) or presence of seminal plasma extracellular vesicles (EVs). Sperm motility and kinematic parameters were evaluated by computer-assisted sperm analysis (CASA). The experiment included three independent biological replicates, each consisting of a pooled semen sample generated from ejaculates of three rams. Values are presented as mean ± SEM calculated from the three biological replicates. Statistical comparisons were performed using linear mixed-effects models (LMMs) with treatment as a fixed effect and biological replicate as a random effect. Full statistical results, including estimates, 95% confidence intervals, and exact p-values, are provided in **Supplementary Table 1**.

| Group | <u>Total motility (%)</u> | <u>Progressive motility (%)</u> | VCL ($\mu\text{m/s}$) | VAP ($\mu\text{m/s}$) | VSL ($\mu\text{m/s}$) | LIN (%) | STR (%) | WOB (%) | ALH ($\mu\text{m}$) | BCF (Hz) |
| --- | --- | --- | --- | --- | --- | --- | --- | --- | --- | --- |
| **Control**<br>(mean ± SEM) | 52.87<br>$\pm 13.57$ | 35.46<br>$\pm 10.35$ | 127.72<br>$\pm 14.59$ | 91.00<br>$\pm 13.99$ | 78.42<br>$\pm 13.88$ | 59.22<br>$\pm 4.90$ | 81.67<br>$\pm 2.58$ | 69.82<br>$\pm 3.72$ | 1.66<br>$\pm 0.12$ | 9.71<br>$\pm 0.35$ |
| **EVs**<br>(mean ± SEM) | 59.99<br>$\pm 20.76$ | 44.29<br>$\pm 22.22$ | 205.49<br>$\pm 87.21$ | 164.51<br>$\pm 74.76$ | 151.23<br>$\pm 72.36$ | 68.80<br>$\pm 4.17$ | 87.65<br>$\pm 2.57$ | 76.82<br>$\pm 2.68$ | 2.37<br>$\pm 0.90$ | 12.52<br>$\pm 1.55$ |

## 4. Discussion

The present study shows that the incorporation of seminal plasma-derived extracellular vesicles (EVs) to semen extender, modulate selected functional responses of ram sperm during cryopreservation, being associated with improved membrane lipid organization, lower intracellular calcium levels, and changes in motility kinematic. In contrast, EV supplementation did not significantly affect sperm viability, oxidative status, acrosomal integrity, or capacitation-associated signaling pathways, including protein tyrosine phosphorylation and PKA activity. These results indicate that their primary action is not on overall cell survival nor on downstream capacitation-related activation. Importantly, these effects were detected after freezing and thawing, indicating that the functional influence of EVs persists despite the substantial membrane destabilization induced by cryopreservation. Collectively, these findings suggest that seminal plasma EVs may act as modulators of sperm membrane physiology by attenuating cryo-induced alterations at the membrane-calcium interface. Cryopreservation-induced disruption of sperm membrane lipid architecture is a well-established phenomenon in ram sperm and has been associated with increased membrane fluidity, loss of lipid asymmetry, and premature capacitation-like changes after thawing (Bailey et al. 2000; Watson 2000; Martínez-Pastor et al. 2010). Previous work from our group demonstrated that cryopreservation destabilizes interactions between seminal plasma proteins and the sperm surface, leading to partial loss of membrane-associated components involved in maintaining lipid organization and functional stability (Ramírez-Vasquez et al., 2019).

In the present study, EV supplementation significantly reduced membrane lipid disorder in post-thaw spermatozoa. The lower Merocyanine 540 fluorescence observed in EV-treated samples suggests improved membrane lipid packing and stability during cryopreservation. This effect may result from direct interactions between EV membranes and the sperm plasma membrane, facilitating lipid exchange or reinforcing membrane structure, as previously proposed in other mammalian species (Sullivan and Saez 2013; Aalberts et al. 2014). In support of this mechanism, recent evidence from our laboratory showed that in ram spermatozoa extracellular vesicles are incorporated in a patchy distribution pattern on the sperm surface, suggesting a direct interaction with and integration into the plasma membrane (Armani et al. 2026). Consistent with this hypothesis, studies in human sperm have shown that seminal plasma exosomes can promote cholesterol and lipid transfer to the sperm membrane, thereby stabilizing membrane architecture and delaying capacitation-associated events (Bechoua et al. 2011). Similar lipid-mediated stabilizing effects have been reported in boar spermatozoa (Piehl et al. 2013). Thus, EVs present during cryopreservation may contribute to maintaining membrane organization after thawing, although the underlying mechanisms remain to be established.

Notably, preservation of membrane organization in EV-treated sperm was associated with reduced intracellular Ca²⁺ levels after thawing. Dysregulation of Ca²⁺ homeostasis is a hallmark of cryodamage and is closely linked to membrane destabilization and premature activation of capacitation-related pathways (Bailey et al. 2000; Martínez-Pastor et al. 2010). Therefore, lower intracellular Ca²⁺ levels may reflect better preservation of membrane barrier function and reduced uncontrolled Ca²⁺ influx. Although EV supplementation affected early capacitation-associated events such as membrane lipid disorder and Ca²⁺ influx, it did not significantly modify downstream signaling pathways, including PKA activation and protein tyrosine phosphorylation, nor did it alter sperm viability, acrosomal status, or oxidative stress. This suggests that EVs do not exert a generalized cytoprotective or antioxidant effect, but rather a targeted action on membrane-associated processes. While EVs have been reported to contain antioxidant enzymes and redox-related proteins (Vojtech et al. 2014; Simon et al. 2018b), ROS levels were unchanged under the experimental conditions of this study, indicating that oxidative stress was not a primary target of EV action. Importantly, acrosomal integrity was also unaffected, suggesting that EVs do not induce premature acrosomal destabilization. Similar observations have been reported in boar spermatozoa, where EVs modulate membrane properties without triggering acrosome reaction (Piehl et al. 2013). Collectively, these findings support the notion that EVs modulate membrane-associated processes during cryopreservation without inducing detectable capacitation-like activation under the conditions tested.

Regarding motility, EV supplementation selectively modified sperm kinematic parameters after thawing without affecting total or progressive motility percentages. In particular, velocity-related parameters, including VCL, VAP, and VSL, were increased in EV-treated spermatozoa, together with higher linearity (LIN), straightness (STR), wobble (WOB), amplitude of lateral head displacement (ALH), and beat-cross frequency (BCF). The simultaneous increase in LIN and STR together with higher velocities indicates that EV supplementation promotes a more linear and progressive movement pattern. This kinematic profile differs from the classical hyperactivated phenotype, which is characterized by high ALH and VCL but low LIN and STR, resulting in highly circular or star-spin trajectories (Suarez and Ho, 2003; Amann and Waberski, 2014). The observed increases in ALH and BCF are of particular interest. ALH reflects the amplitude of flagellar bending, while BCF indicates the frequency of flagellar beat cycles (Mortimer, 1997). The concurrent elevation of both parameters suggests that EV supplementation enhanced flagellar activity and energy expenditure in motile spermatozoa. This is consistent with our previous observations of lower intracellular calcium levels in EV-treated samples, as calcium signaling is a key regulator of flagellar beat frequency and amplitude (Suarez, 2008; Publicover et al., 2008). Whether this increased flagellar activity translates into improved sperm transport through the female reproductive tract or enhanced fertilization competence remains to be determined. Importantly, the observed kinematic changes cannot be directly interpreted as evidence of improved fertilizing ability, as no functional assays were performed in this study. While the more linear and active movement pattern could be advantageous for sperm progression through the oviductal environment (Holt and Fazeli, 2010), this interpretation remains speculative. Future studies incorporating sperm-oocyte interaction assays and fertility assessments will be necessary to determine whether the kinematic and membrane-related changes reported here translate into improved reproductive outcomes.

Although EV supplementation was associated with improved membrane lipid organization, lower intracellular Ca²⁺ levels and changes in sperm motility kinematics, these functional changes should not be directly interpreted as evidence of enhanced fertilizing ability. To address this, post-thawing sperm functional parameters can be considered a laboratory supported contribution, however sperm-oocyte interaction assays, in vitro fertilization, or in vivo fertility assessments are needed. Furthermore, it should be noted that the experimental design of this work was based on pooled ejaculates and three independent biological replicates. While this approach minimized individual ejaculate viability and provided sufficient biological material for EV isolation, characterization, and comprehensive functional analyses, it did not allow for the assessment of individual ram responses to EV supplementation. Therefore, future studies evaluating reproductive outcomes will be necessary to determine whether the membrane- and calcium-associated changes reported here translate into improved fertility. The present findings are consistent with previous reports in bovine and porcine species showing beneficial effects of EVs on sperm function and cryotolerance (Lange-Consiglio et al. 2022; Esmaeili et al. 2025). However, to our knowledge, this is the first demonstration of a functional role for seminal plasma-derived EVs during semen cryopreservation in rams. Given the high sensitivity of ovine sperm to cryoinjury, EV supplementation represents a promising biological approach to modulate selected sperm functional parameters during cryopreservation. However, additional studies evaluating fertilization and reproductive outcomes will be required to determine its practical value for improving cryopreservation protocols.

## 5. Conclusions

In conclusion, despite the limited number of biological replicates, extender supplementation with seminal plasma-derived extracellular vesicles for ram sperm cryopreservation was associated with improved membrane lipid organization, lower intracellular calcium levels, and changes in post-thaw motility kinematics, while maintaining sperm viability. No effect was observed on sperm, oxidative status, acrosomal integrity, or capacitation-associated signaling under the conditions evaluated. These findings provide initial evidence supporting the potential use of seminal plasma-derived EVs as biological supplement during ram semen cryopreservation. However, further studies evaluating fertilizing ability and reproductive outcomes will be necessary to determine the functional significance of the observed effects.

## Author contributions

AN and AC contributed to the conceptualization of the study, supervision, and project administration. AN, TA, LZ, MBA, FH, and AC contributed to the design and development of the methodology. AN and TA analyzed the results. AN, TA, LZ, and AC wrote the manuscript. All co-authors were involved in revising the manuscript.

---

**Supplementary Figure 1.**
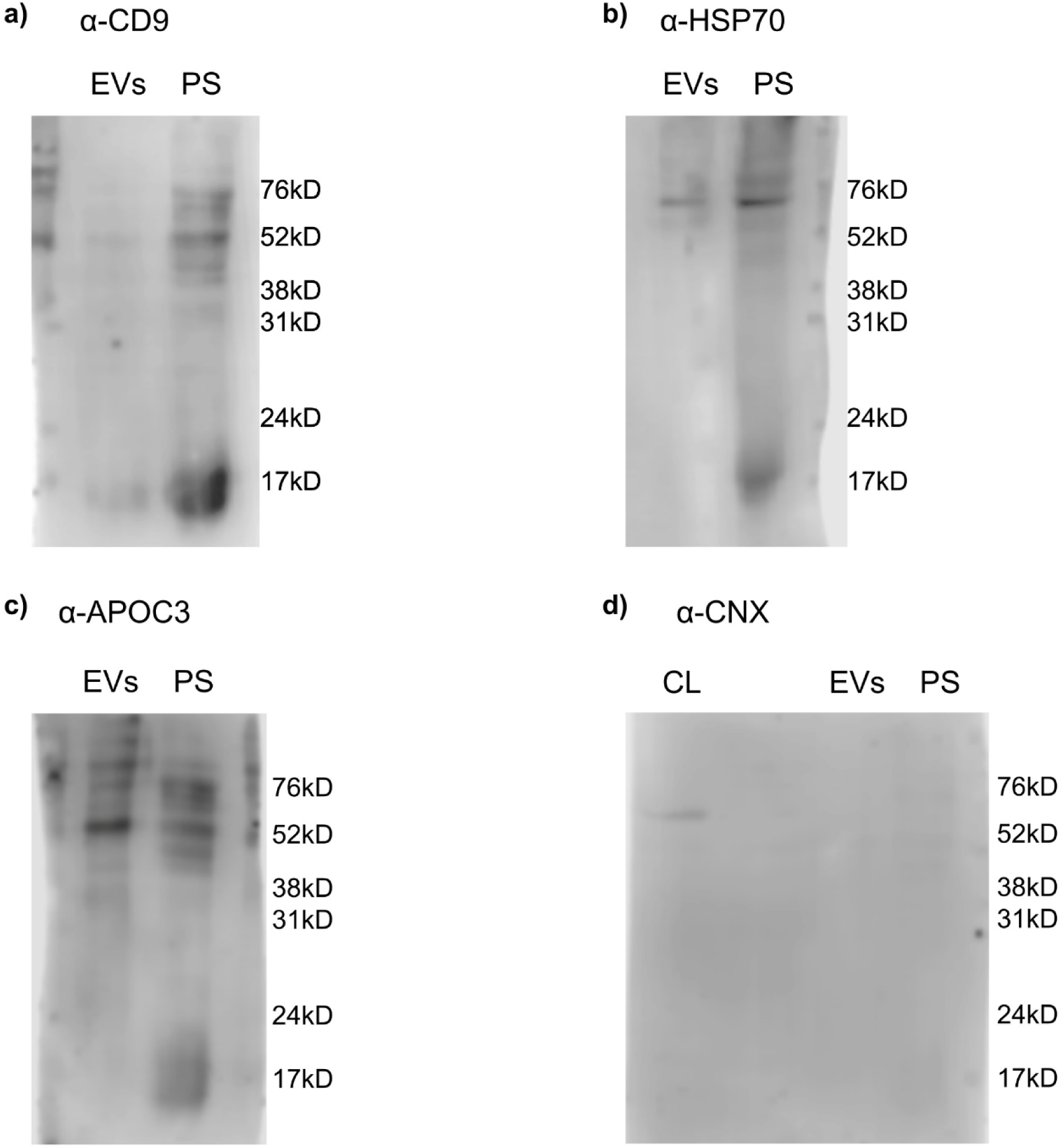
Full-length Western blots used for the characterization of ram seminal plasma extracellular vesicles. Representative immunoblots showing detection of (a) CD9, (b) HSP70, (c) ApoC3, and (d) calnexin (CNX) obtained from three independent pooled seminal plasma samples, each prepared from ejaculates of three rams.

**Supplementary Figure 2.**
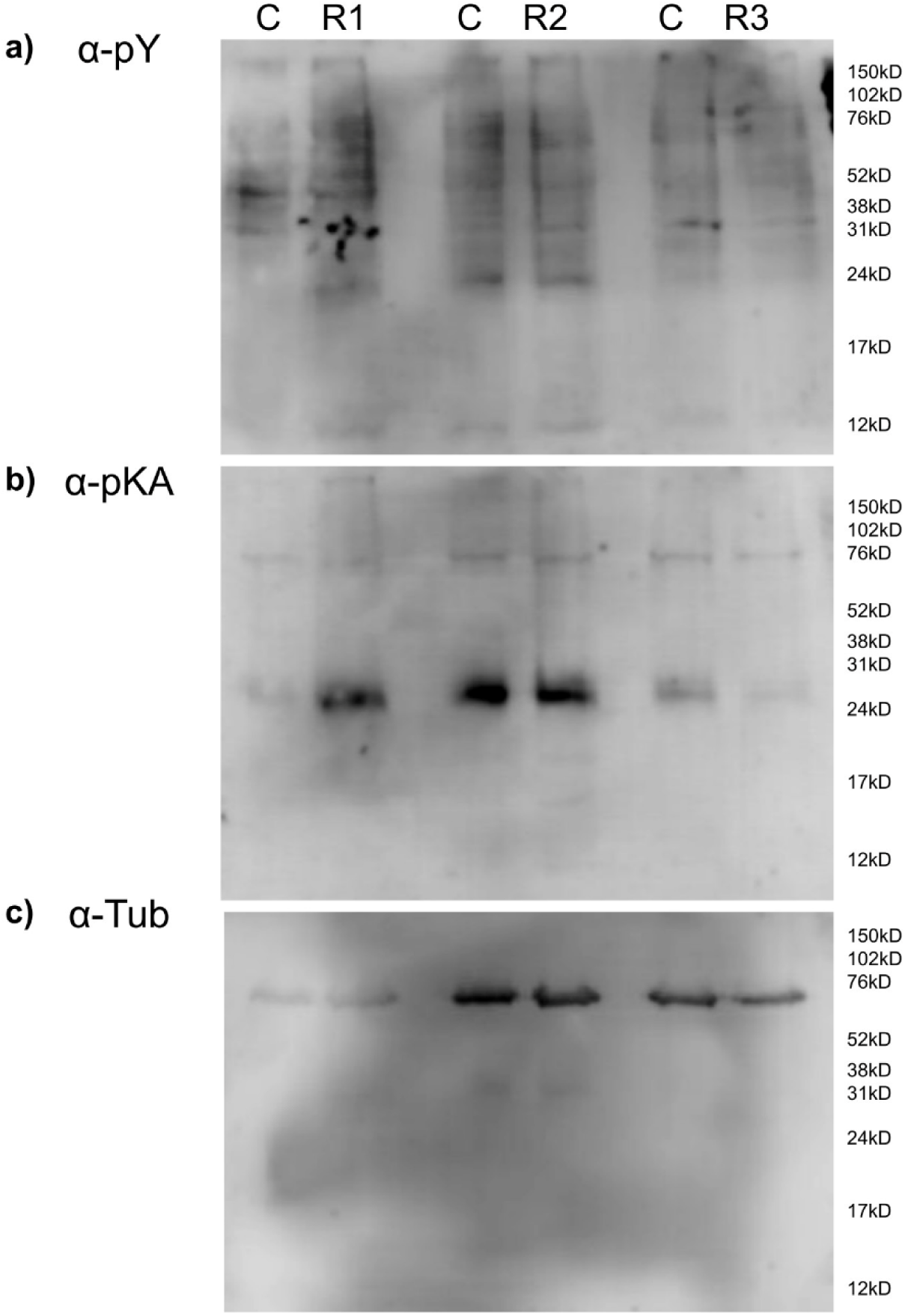
Full-length Western blots corresponding to the protein phosphorylation analyses shown in **Figure 3**. Representative immunoblots obtained from three independent pooled semen samples, each prepared from ejaculates of three rams (R1–R3), using antibodies against (a) phosphotyrosine (α-pY), (b) PKA substrates (α-PKA), and (c) α-tubulin (α-Tub). For each biological replicate, Control (C) and EV-treated samples are shown. Molecular weight markers (kDa) are indicated on the right. These uncropped blots were used for the densitometric analyses presented in **Figure 3**.

**Supplementary Table 1.** Summary of statistical analyses for all evaluated parameters. Data were analyzed using linear mixed-effects models (LMMs) or generalized linear mixed-effects models (GLMMs) with treatment (EVs vs. Control) as a fixed effect and biological replicate as a random effect. For LMMs, the treatment effect estimate represents the difference between EV and Control groups (EV - Control). For the GLMM beta model (M540), the estimate is presented as an odds ratio (OR) with its 95% confidence interval. Negative estimates indicate lower values in the EV group compared to Control, while positive estimates indicate higher values in the EV group. Significant effects (p < 0.05) are highlighted in bold.

| Variable | Model | Treatment effect estimate<br>(EV vs Control) | 95% CI | p |
| --- | --- | --- | --- | --- |
| Viability | LMM | 7.34 | -7.56 to 22.2 | 0.215 |
| PNA | LMM | -7.19 | -27.5 to 13.2 | 0.343 |
| ROS | LMM | 0.49 | -0.09 to 1.07 | 0.075 |
| Calcium | LMM | -5.63 | -11.3 to 0.008 | 0.048 |
| M540 | GLMM beta | OR=0.187 | 0.109 to 0.322 | <0.0001 |
| Total motility | LMM | 7.12 | -0.74 to 15.0 | 0.068 |
| Progressive motility | LMM | 8.83 | -5.25 to 22.9 | 0.176 |
| pY phosphorylation | LMM | -4.76 | -12.6 to 3.11 | 0.168 |
| PKA phosphorylation | LMM | 9.64 | -5.06 to 24.3 | 0.153 |
| VCL | LMM | 84.3 | 69.2 to 99.4 | <0.0001 |
| VAP | LMM | 78.7 | 63.6 to 93.7 | <0.0001 |
| VSL | LMM | 77.6 | 62.2 to 92.9 | <0.0001 |
| LIN | LMM | 9.15 | 6.82 to 11.5 | <0.0001 |
| STR | LMM | 5.79 | 4.06 to 7.53 | <0.0001 |
| WOB | LMM | 6.63 | 4.88 to 8.38 | <0.0001 |
| ALH | LMM | 0.783 | 0.615 to 0.951 | <0.0001 |
| BCF | LMM | 2.6 | 1.93 to 3.27 | <0.0001 |

